# High Transcriptional Activity and Diverse Functional Repertoires of Hundreds of Giant Viruses in a Coastal Marine System

**DOI:** 10.1101/2021.03.08.434518

**Authors:** Anh D Ha, Mohammad Moniruzzaman, Frank O Aylward

## Abstract

Viruses belonging to the *Nucleocytoviricota* phylum are globally-distributed and include members with notably large genomes and complex functional repertoires. Recent studies have shown that these viruses are particularly diverse and abundant in marine systems, but the magnitude of actively replicating *Nucleocytoviricota* present in ocean habitats remains unclear. In this study, we compiled a curated database of 2,431 *Nucleocytoviricota* genomes and used it to examine the gene expression of these viruses in a 2.5-day metatranscriptomic time-series from surface waters of the California Current. We identified 145 viral genomes with high levels of gene expression, including 90 *Imitervirales* and 49 *Algavirales* viruses. In addition to recovering high expression of core genes involved in information processing that are commonly expressed during viral infection, we also identified transcripts of diverse viral metabolic genes from pathways such as glycolysis, the TCA cycle, and the pentose phosphate pathway, suggesting that virus-mediated reprogramming of central carbon metabolism is common in oceanic surface waters. Surprisingly, we also identified viral transcripts with homology to actin, myosin, and kinesin domains, suggesting that viruses may use them to manipulate host cytoskeletal dynamics during infection. We performed phylogenetic analysis on the virus-encoded myosin and kinesin proteins, which demonstrated that most belong to deep-branching viral clades, but that others appear to have been acquired from eukaryotes more recently. Our results highlight a remarkable diversity of active *Nucleocytoviricota* in a coastal marine system and underscore the complex functional repertoires expressed by these viruses during infection.

**Importance:** The discovery of giant viruses has transformed our understanding of viral complexity. Although viruses have traditionally been viewed as filterable infectious agents that lack metabolism, giant viruses can reach sizes rivalling cellular lineages and possess genomes encoding central metabolic processes. Recent studies have shown that giant viruses are widespread in aquatic systems, but the activity of these viruses and the extent to which they reprogram host physiology *in situ* remains unclear. Here we show that numerous giant viruses consistently express central metabolic enzymes in a coastal marine system, including components of glycolysis, the TCA cycle, and other pathways involved in nutrient homeostasis. Moreover, we found expression of several viral-encoded actin, myosin, and kinesin genes, indicating viral manipulation of the host cytoskeleton during infection. Our study reveals a high activity of giant viruses in a coastal marine system and indicates they are a diverse and underappreciated component of microbial diversity in the ocean.

## Introduction

Large dsDNA viruses of the phylum *Nucleocytoviricota*, commonly referred to as “giant viruses”, are a diverse group of double-stranded DNA viruses with virion sizes reaching 1.5 μm, comparable to the sizes of many cellular lineages (1–4). A recently-proposed taxonomy of this viral phylum delineated 6 orders and 32 families (5), including the previously-established families *Asfarviridae, Poxviridae, Marseilleviridae, Iridoviridae, Phycodnaviridae*, and *Mimiviridae*. Members of the *Nucleocytoviricota* are known to infect a broad range of eukaryotic hosts; members of the *Poxviridae* and *Iridoviridae* families infect numerous metazoans, members of the *Imitervirales* and *Algavirales* orders infect a wide range of algae and other protists, and members of the *Asfuvirales* infect a mixture of metazoan and protist hosts (6–10). Members of the *Nucleocytoviricota* typically harbor exceptionally large genomes that are often > 300 kbp in length, and in some cases as large as 2.5 Mbp (9). Numerous studies have noted the unusually complex genomic repertoires of viruses in this group that include many genes typically found only in cellular lineages, such as those involved in the TCA cycle (11–13), glycolysis (12), amino acid metabolism (14), light sensing (15, 16), sphingolipid biosynthesis (17, 18), eukaryotic cytoskeleton (19), and fermentation (19). These complex metabolic repertoires are thought to play a role in the manipulation of host physiology during infection, in effect transforming healthy cells into reprogrammed “virocells” that more efficiently produce viral progeny (12, 20, 21).

The genomic complexity of *Nucleocytoviricota* together with reports that they are abundant in global marine environments has raised questions regarding the extent to which they manipulate the physiology of their hosts and thereby influence global biogeochemical cycles (22). Cultivated representatives of these viruses are known to infect a broad array of ecologically-important eukaryotic algae, including dinoflagellates, haptophytes, brown algae, and chlorophytes, among others (23–26), which suggests that they could play important roles in shaping the activity and composition of the microbial eukaryotic community. Even so, our understanding of the distribution and activity of *Nucleocytoviricota* in marine systems has lagged behind that of other viral groups owing to their large size, which precludes their presence in the small size fractions typically analyzed in viral diversity studies (10). Some early studies were prescient in suggesting that these viruses are a diverse and underappreciated component of viral diversity in the ocean (27–30). More recent work has confirmed this view and revealed that a diverse range of *Nucleocytoviricota* inhabit the global ocean and likely contribute to key processes such as algal bloom termination and carbon export (12, 31–36). Some metatranscriptomic studies have also begun to note the presence of *Nucleocytoviricota* transcripts in marine samples, confirming their activity (37–39). Although these studies have vastly expanded our knowledge of the diversity and host range of *Nucleocytoviricota* in the ocean, the activity of these viruses and the extent to which they use cellular metabolic genes they encode during infection remains unclear.

In this study, we constructed a database of 2,436 annotated *Nucleocytoviricota* genomes for the purpose of evaluating the gene expression landscape of these viruses. We then leveraged a previously published metatranscriptome time-series from surface waters of the California Current (39) to assess the daily activity of *Nucleocytoviricota* in this environment. We show that hundreds of these viruses, primarily of the orders *Imitervirales* (including the family *Mimiviridae*) and *Algavirales* (including the families *Phycodnaviridae* and *Prasinoviridae*) were consistently active during this period and frequently expressed genes involved in central carbon metabolism, light harvesting, oxidative stress reduction, and lipid metabolism. Unexpectedly, we also found expression of several viral-encoded cytoskeleton genes, including those that encode the motor proteins myosin and kinesin, and we performed a phylogenetic analysis demonstrating that *Nucleocytoviricota* commonly encode deep-branching enzymes in these protein families. Our findings highlight the surprisingly high activity of *Nucleocytoviricota* in marine systems and suggest they are an underappreciated component of viral diversity in the ocean.

## Results and Discussion

We examined a metatranscriptomic dataset of 16 timepoints sampled over a 60-hour period that were obtained from microbial communities previously reported for the California current system (39). This dataset is ideal for examining the activity of marine *Nucleocytoviricota* because it targeted the >5 μm size fraction, which is enriched in many of the eukaryotic plankton that these viruses are known to infect. Altogether, we identified 145 *Nucleocytoviricota* genomes with metatranscriptomic reads mapping over the sampling period, including 90 *Imitervirales*, 49 *Algavirales*, 4 *Pandoravirales*, 1 *Pimascovirales*, and 1 *Asfuvirales* (Fig 1, see Methods for details). Expression levels of the 145 genomes with reads mapping are summarized in Supplemental Data S1. Of the 145 genomes detected, 9 are complete genomes of cultivated viruses, while 136 are genomes derived from cultivation-independent methods (i.e., metagenome-assembled genomes, or MAGs). Almost half (66) of the 145 *Nucleocytoviricota* genomes have genome or assembly sizes > 300 kbp (Fig. S1), indicating that a large fraction of the active viruses have large genomes with complex functional repertoires. The abundance of each *Nucleocytoviricota* genome in the transcriptomes was positively correlated to the number of genes that could be recovered for that virus (Fig. 1a), which is expected given that only highly-expressed viral genes will be recovered for low-abundance viruses in the dataset. Over 80% of the predicted genes in five genomes were recovered throughout the sampling period (one *Imitervirales* and four *Algavirales* viruses), indicating that the recovery of most transcripts is possible for some abundant viruses (Fig 1).

**Figure 1.**
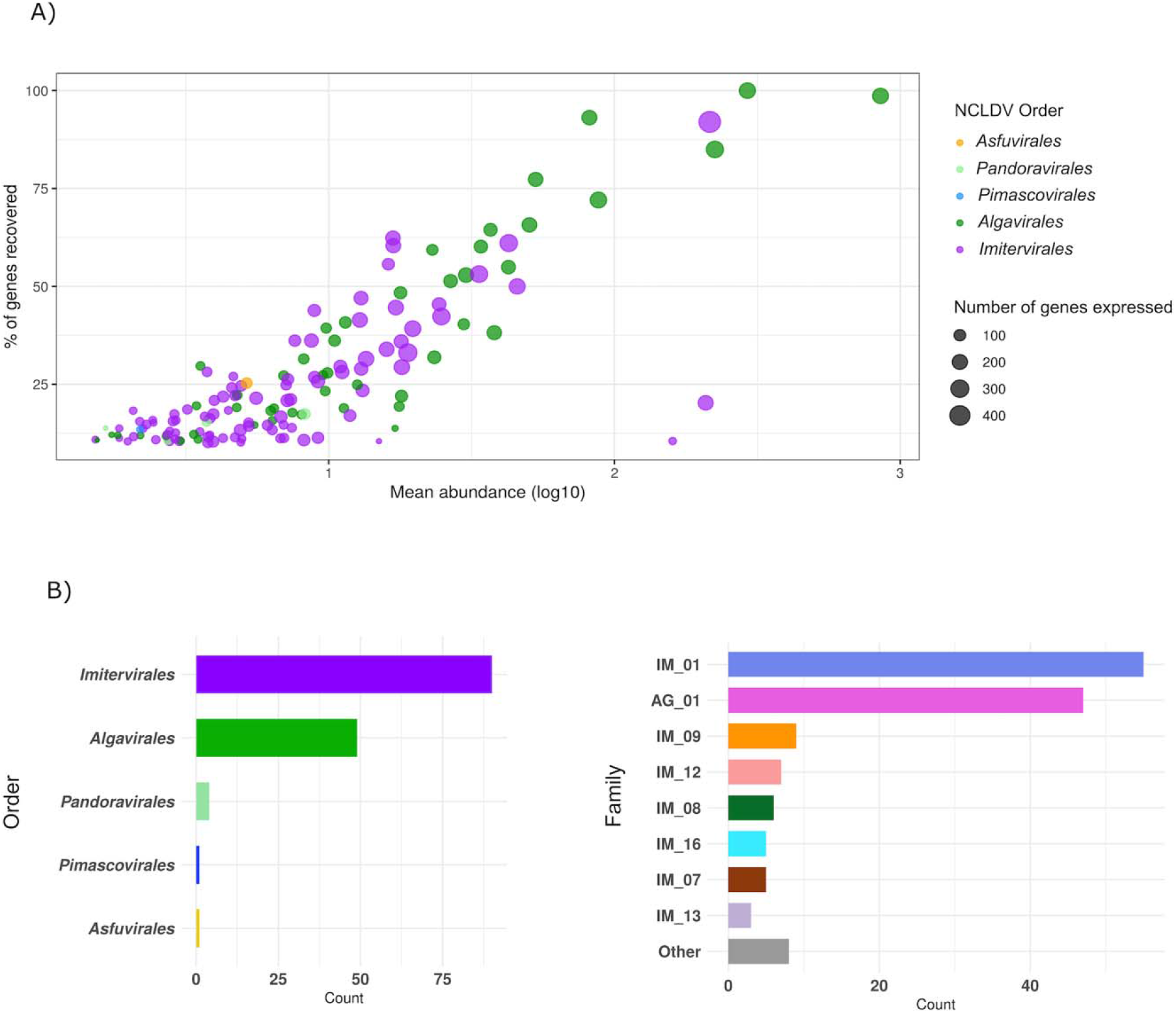
(A) General statistics of the 145 *Nucleocytoviricota* genomes. Each dot represents a viral genome, the x-axis shows the mean abundance of the genome across 16 timepoints (log-transformed TPM), the y-axis shows the percentage of genes in a viral genome that were recovered across all samples, and the dot size is scaled to the total number of genes recovered. (B) The taxonomic distribution of the 145 genomes. The x-axis represents the total number of viruses in each taxon, and the y-axis shows the *Nucleocytoviricota* order (left) and family (right). In the clade plot (right), families with less than three members are collapsed down into “Other” category. Abbreviations: IM: *Imitervirales*, AG: *Algavirales*.

We constructed a multi-locus phylogenetic tree of the 145 viruses present in the transcriptomes together with 1,458 references so that we could examine the phylogenetic distribution of the active *Nucleocytoviricota* (Fig 2). The topology of the resulting tree is consistent with a previous tree we constructed using similar methods (12). Given the large diversity of *Imitervirales* and *Algavirales* viruses in the dataset, we also analyzed the family-level diversity within these orders, using the recently-proposed taxonomic framework for the *Nucleocytoviricota* as a guide (5). Of the 145 viruses that were abundant in the metatranscriptomes, the *Mesomimiviridae (Imitervirales* family 1) and *Prasinoviridae (Algavirales* family 1) were by far the most well-represented (55 and 47 genomes, respectively, Fig 1B). Interestingly, both of these families also include cultivated representatives that are known to infect marine protists, which suggests possible hosts for the viruses in the transcriptomes. The *Mesomimiviridae* contains haptophyte viruses that infect the genera *Phaeocystis* and *Chrysochromulina (40–42)*, while the *Prasinoviridae* contains Prasinoviruses that are known to infect *Ostreococcus, Bathycoccus*, and *Micromonas (8)*. Transcripts mapping to isolate *Ostreococcus* viruses within the *Prasinoviridae* was previously reported in the original analysis of these metatranscriptomes (39); in addition to recovering these genomes, we also identified transcripts mapping to an additional 39 metagenome-derived *Prasinoviridae* genomes, highlighting the diversity of viruses within this clade that were active in the same community during the sampling period. A wide range of other *Imitervirales* viruses were also present in addition to the *Mesomimiviridae* (Fig 1c): *Mimiviridae* (IM_16), which includes the original *Acanthamoeba polyphaga* Mimivirus and its close relatives, IM_12 containing the recently-cultivated Tetraselmis virus (TetV) (19), IM_08, which includes the recently-reported Choanoflagellate virus (43), IM_09, which includes *Aureococcus anophagefferens* virus (AaV) (26), and *Imitervirales* families 7 and 13 which contain no cultivated representatives. One *Pimascovirales* virus (GVMAG-M-3300009467-15) and one *Asfuvirales* virus (GVMAG-M-3300027833-19) were also abundant in the transcriptomes, suggesting they may also infect microbial eukaryotes in the community. The *Asfuvirales* virus found here was also recently identified in waters off the coast of South Africa (7), indicating that it may be broadly distributed in the ocean. The presence of numerous divergent families of *Nucleocytoviricota* in the metatranscriptomes supports the view that marine systems harbor an immense diversity of these viruses that are actively infecting a wide variety of protists hosts.

**Figure 2.**
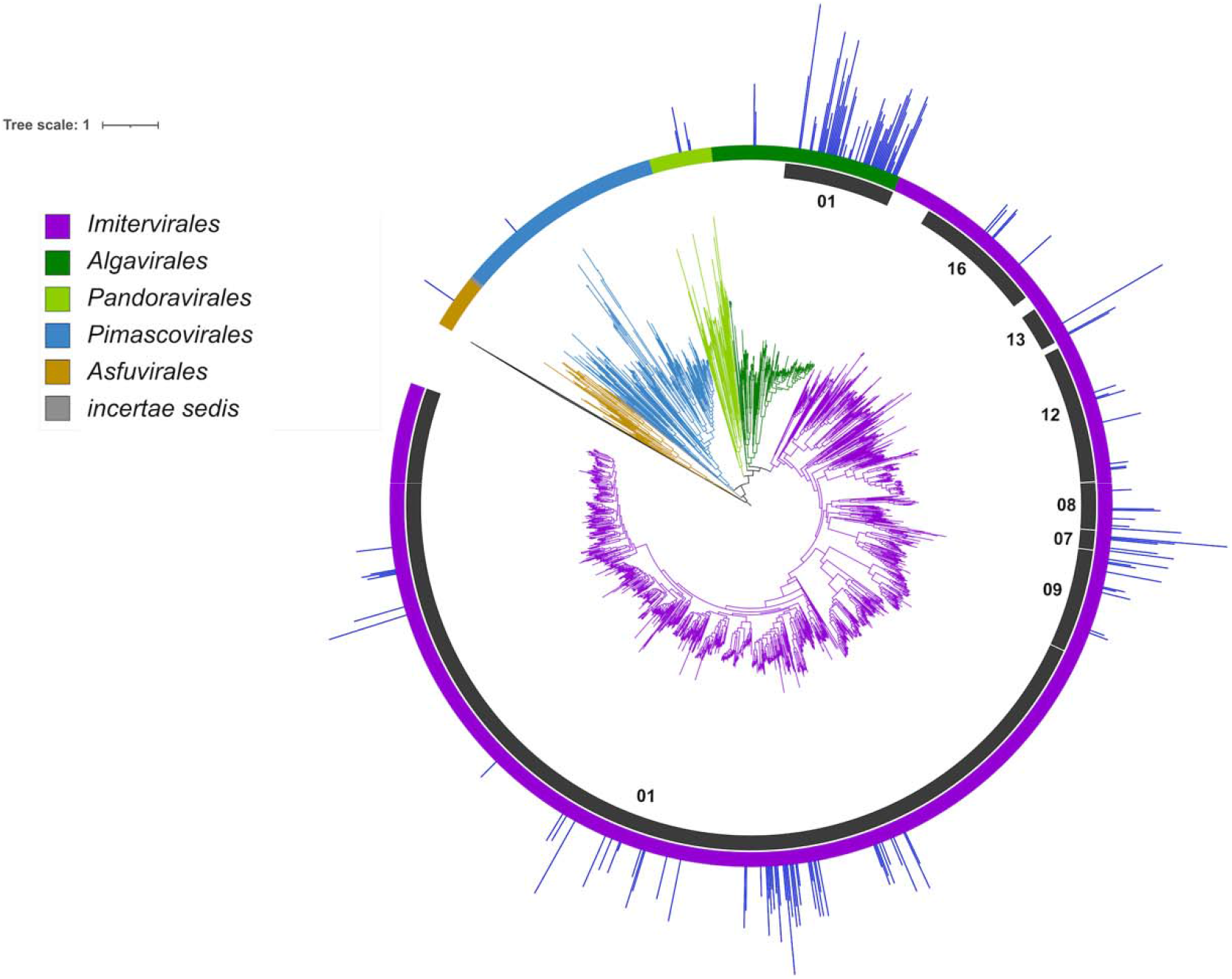
Phylogeny of *Nucleocytoviricota* with mean read mapping (TPM, log10-transformed) on the outermost bar plot. The order-level classification of *Nucleocytoviricota* is denoted by the branch colors and the outer color strip, and particular families of interest are highlighted in the inner ring. IM_01 corresponds to the *Mesomimiviridae*, IM_16 corresponds to the *Mimiviridae*, and AG_01 corresponds to the *Prasinoviridae*.

We examined summed whole-genome transcriptional activity of the 145 *Nucleocytoviricota* to identify potential diel cycles of viral activity. We found 6 viruses that had significantly higher expression during nighttime periods (Fig 3A, Mann-Whitney U test, corrected p-value < 0.1). These transcriptional patterns were not strictly diel because their peak of expression was found at different times during the night, but they nonetheless suggest that viral activity may be higher at night for some of these viruses. We also analyzed whole-genome transcriptional activity using a weighted network-based approach that groups viruses with similar temporal transcriptional profiles into modules. This analysis revealed two primary modules, one of which contained 60 viruses and had a module-wide transcriptional profile with notable peaks at nighttime periods (Module 1, Fig 3B). We found no significant diel cycles in the *Nucleocytoviricota* whole genome transcriptional profiles when we tested for strict diel periodicity (RAIN, p-value < 0.1, see Methods), suggesting that, although viral activity appears higher at night for some viruses, pronounced and significant diel cycling of viral activity was not detectable. This does not necessarily suggest that diel cycles are uncommon in marine *Nucleocytoviricota*; previous studies have found that the detection of diel cycles in metatranscriptomic data can require longer time-series (>20 time-points) and may only be feasible for the most abundant community members for which gene expression is high (44).

**Figure 3.**
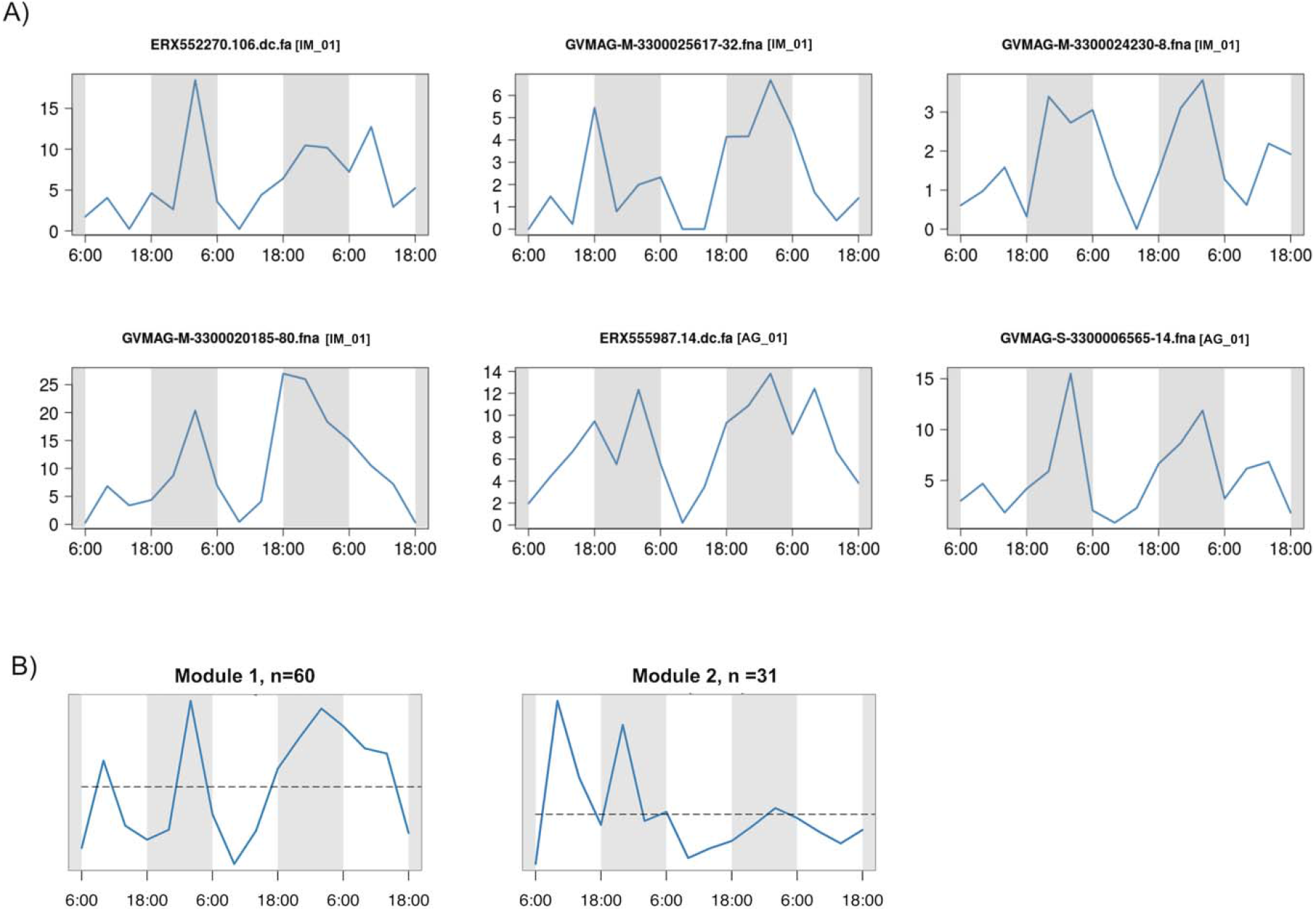
Expression level of *Nucleocytoviricota* during the 16 timepoints studied (A) Whole-genome expression profiles of six genomes with significant difference in expression level between daytime and nighttime (Mann-Whitney U test p< 0.1). The x-axis represents the 16 timepoints and the y-axis represents the total expression (TPM). *Nucleocytoviricota* family classifications are shown in brackets. Abbreviations: IM: *Imitervirales*, AG: *Algavirales*. (B) Eigengene of the genome co-expression modules. The x-axis represents the 16 timepoints and the y-axis represents the eigengene expression values.

The general trend of higher night-time expression of *Nucleocytoviricota* is consistent with previous studies of marine viruses. A culture-based study of *Ostreococcus tauri* virus 5 found that viral transcripts increased markedly at night (45). Two of the *Nucleocytoviricota* with higher nighttime expression in our analysis belong to the *Prasinoviridae* (AG_01), which includes cultivated viruses that infect *Ostreococcus* and other prasinophytes, suggesting that higher nighttime activity may therefore be a common trait in this family. Previous work on bacteriophage has revealed peak viral transcription at various times throughout a 24-hr period depending on the viral group (46–48), although in many cases gene expression was found to peak near dusk. It has been hypothesized that high viral expression at dusk is linked to the energetic state of phototrophic host cells such as *Prochlorococcus*, which grow throughout the day and divide at dusk. Viruses active near the end of the host growth period may therefore have more cellular resources to exploit for virion production (46). Whether or not a similar dynamic is at play with *Nucleocytoviricota* is unclear; the *Prasinoviridae* gene expression appears to peak after dusk, but this may be caused in part by a longer infection program of these viruses compared to cyanophages. The four *Mesomimiviridae* with diel transcriptional signatures that peak in the evening (IM_01, Fig. 3) may also infect phototrophic hosts where similar factors are at play. Many of the *Nucleocytoviricota* we detected in the transcriptomes likely infect heterotrophic hosts, however, and it remains unclear if there would be reason to expect a diel infection pattern in these viruses. Further work is therefore needed to examine the potential role of diel cycling in these viruses in more detail.

Throughout the sampling period, we detected expression of numerous viral genes involved in diverse metabolic processes (Fig 4, Fig S2, Supplemental Data S2). As expected, we found high expression of viral core genes, including the major capsid protein, family B DNA Polymerase, A32 packaging enzyme, VLTF3 transcriptional factor, and superfamily II helicase (Fig. 4); this is consistent with both culture-based and culture-independent studies of *Nucleocytoviricota* that have found most of these genes, and in particular the capsid proteins, to be highly expressed during infection (45, 49–52). Chaperones from the Hsp70 and Hsp90 families were also highly expressed in numerous *Nucleocytoviricota*, consistent with some previous transcriptome studies (50, 52); these likely play a role in protein folding in virus factories, particularly for capsids (3, 52). We also detected transcripts involved in mitigating cellular stress, such as glutathione-S-transferase (GST) and superoxide dismutase (SOD). SOD is involved in the detoxification of reactive oxygen species produced by hosts as a defense mechanism during infection, and it has been postulated that viral-encoded SODs may allow some viruses to infect a broader array of hosts (53). GST may be involved in the detoxification of electrophilic metabolites, and the host version of this gene was also found to be overexpressed in several time-points of an infection experiment of *Aureococcus anophagefferens* virus and its host (54).

**Figure 4.**
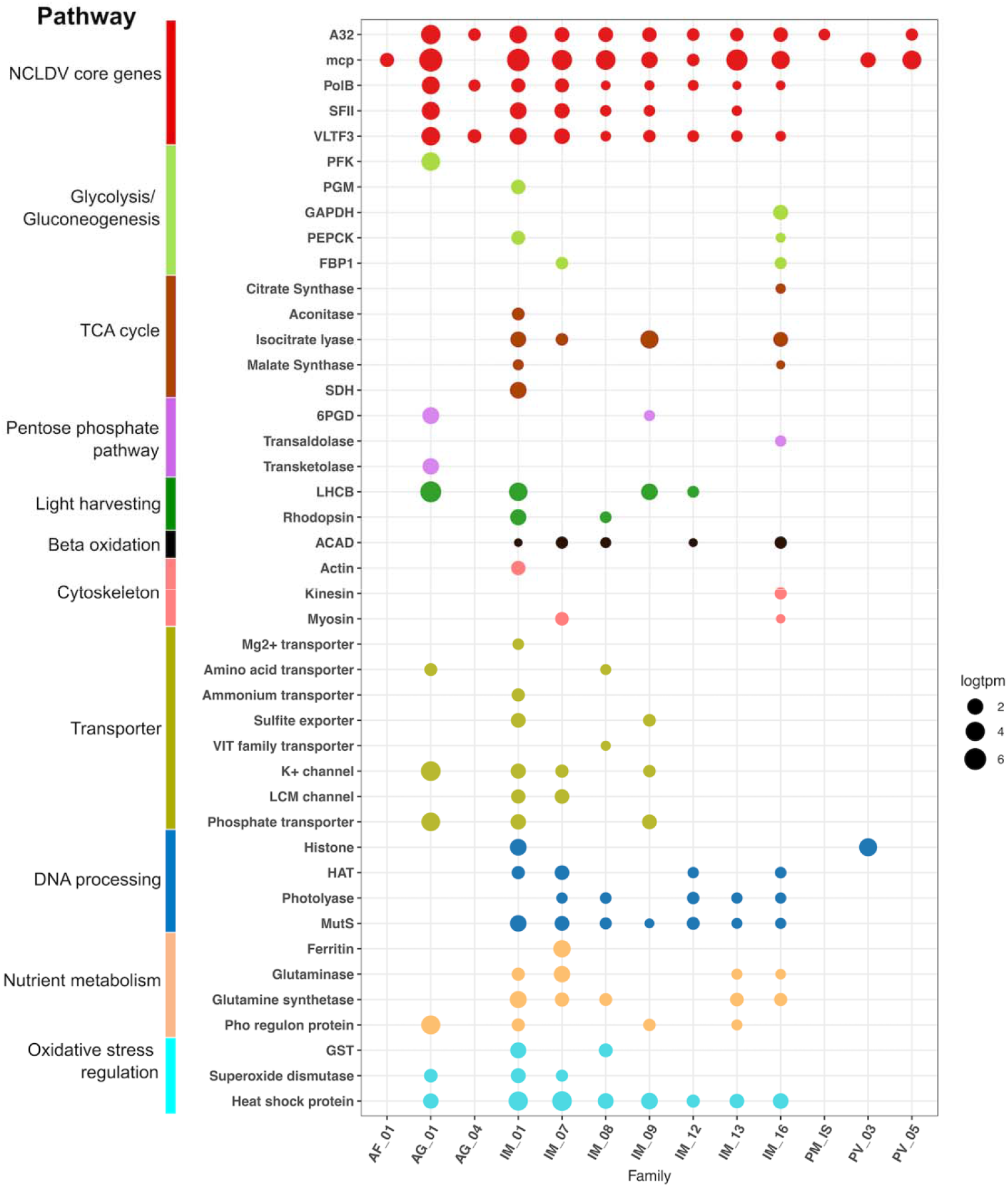
Metabolic genes expressed in the transcriptomes. The x-axis shows different viral clades and the y-axis denotes the functional annotation. The sizes of the bubbles represent the total abundance of the gene in TPM (log-transformed) and the colors show the functional category. Abbreviations: a) Family assignments. AF: *Asfuvirales*, IM: *Imitervirales*, AG: *Algavirales*, PM: *Pimascovirales*, PV: *Pandoravirales*, IS: *incertae sedis*. b) Gene function. PFK: Phosphofructokinase, PGM: Phosphoglycerate mutase, GAPDH: Glyceraldehyde 3-P dehydrogenase, PEPCK: Phosphoenolpyruvate carboxykinase, FBP1: Fructose-1-6-bisphosphatase, SDH: Succinate dehydrogenase, 6PGD: 6-phosphogluconate dehydrogenase, LHCB: Chlorophyll a/b binding protein, ACAD: Acyl-CoA dehydrogenase, HAT: Histone acetyltransferase, GST: Glutathione S-transferase.

We also detected numerous transporters predicted to target nitrogen, sulfur, and phosphorus-containing nutrients, consistent with the view that *Nucleocytoviricota* manipulate the nutrient environment of their hosts during infection. Among these, we identified expression of several viral ammonium transporters; previous work with an ammonium transporter encoded by *Ostreococcus tauri virus* 6 has shown that this gene is expressed during infection and manipulates nutrient uptake by the host cell (55). Viral-encoded phosphate transporters are also common in many marine *Nucleocytoviricota* (12, 56), and expression of these genes further emphasizes their likely role in viral-mediated nutrient transport during infection. Other genes involved in nutrient homeostasis were also well-represented in the metatranscriptomes, including ferritin, glutaminase, glutamine synthetase, and Pho regulon components (Fig. 4), consistent with recent findings that these genes are also common in these viruses (12).

Recent work has noted that many *Nucleocytoviricota* encode enzymes involved in central metabolic pathways, and it has been postulated that these have a role in the reprogramming of healthy cells into virocells during infection (12). We detected expression of many of these central metabolic enzymes throughout the sampling period; genes involved in glycolysis, the TCA cycle, the pentose phosphate pathway, and beta oxidation were all present in the metatranscriptomes, and most were consistently expressed across timepoints, especially in the *Imitervirales* (Fig 4, Fig S2). A recent study of TCA cycle enzymes encoded by a *Pandoravirus* isolate has suggested that these enzymes have activity consistent with bioinformatic predictions, and that some are packaged into virions and would therefore influence host physiology soon after initial infection (57). Together with our results, this suggests that viral manipulation of central metabolic processes is a widespread mechanism employed by many *Nucleocytoviricota*, and that this is a common occurrence in viral infections in the ocean. This raises the question of what proportion of the protist community is infected by *Nucleocytoviricota* at any given time, and thereby has an altered metabolic state. Previous studies examining infected hosts have found that 27-37% of cells can be infected by *Nucleocytovicota* during an algal bloom (58, 59); given the possibility that multiple viruses may infect the same host population, the total number of infected cells may be substantially higher. This leads to the surprising possibility that a large proportion of protist populations in some marine systems may consist of “virocells” in which their metabolism is substantially altered by viruses that infect them, at least for some periods of high viral activity.

Translation-related genes were once thought to be trademarks of cellular organisms until the discovery of their presence in viruses of the family *Mimiviridae*, and have since become one of the most noteworthy features of the genomic repertoires of giant viruses. We observed high expressions of several aminoacyl–tRNA synthetase genes (aaRSs), including those that encode asparaginyl-tRNA synthetases, lysyl-tRNA synthetases, tyrosyl-tRNA synthetases, and prolyl-tRNA synthetases. Transcripts mapping to these genes were found from *Imitervirales* genomes in the *Mimiviridae, Mesomimiviridae*, IM_07, IM_12, and IM_13 families. Proteins encoded by the aaRS genes are responsible for the interaction between tRNAs and their amino acids, and have been previously found in diverse members of the *Mimiviridae* (60, 61) and may be associated with an increase of viral fitness (62). We also detected viral transcripts from *Imitervirales* genes with homology to genes involved in translation initiation, such as translation initiation factor 4E (IF4E), translation initiation factor SUI1, translation initiation factor 3 (IF3). Translation elongation factors EF-TU, EIF-5a, and EF-P were also expressed, especially in the *Mimiviridae* and families IM_07, IM_09, and IM_13. This is consistent with the finding that many mimivirus genomes encode various translation factors (63, 64). The expression of viral genes involved in translation suggests a degree of independence from the host’s translational apparatus machinery during active infections occurring in oceanic waters.

Surprisingly, we also found several transcripts matching to predicted cytoskeletal proteins, including actin, myosin, and kinesin. Recent studies have identified divergent actin, myosin, and kinesin homologs in *Nucleocytoviricota*, and in some cases it has been suggested that they played a role in the emergence of this protein in extant eukaryotic lineages (65–67). We found three expressed viral genes with predicted myosin motor domains and one with kinesin domains, and subsequent analysis of all *Nucleocytoviricota* genomes in our reference database recovered an additional 109 myosin and 200 kinesin homologs. To examine if these genes were recently acquired from cellular hosts, we performed a phylogenetic analysis of the viral myosin and kinesin proteins with references; our results indicate that the viral proteins typically form separate branches that are distinct from their eukaryotic homologs, except in a few cases where it appears some *Nucleocytoviricota* have acquired cellular copies more recently (Fig 5). This is consistent with recent studies that have revealed a dynamic gene exchange between *Nucleocytoviricota* and their hosts that in some cases dates back to the early diversification of eukaryotes (16, 68–71). The expression of viral genes with homology to actin, myosin, and kinesin domains suggests that these viruses may use these proteins to manipulate host cytoskeletal dynamics during infection; it is possible that these proteins play a role in the establishment and maintenance of perinuclear viral factories, which are responsible for viral morphogenesis, or for the subcellular trafficking of viral particles. In *Aureococcus anophagefferens* cytoskeletal genes are downregulated during infection by it’s virus AaV (54); if this is common across other host-virus systems it would suggest that *Nucleocytoviricota*-encoded myosin, kinesin, and actin proteins may help to maintain the functioning of the cytoskeleton during later periods of infection.

**Figure 5.**
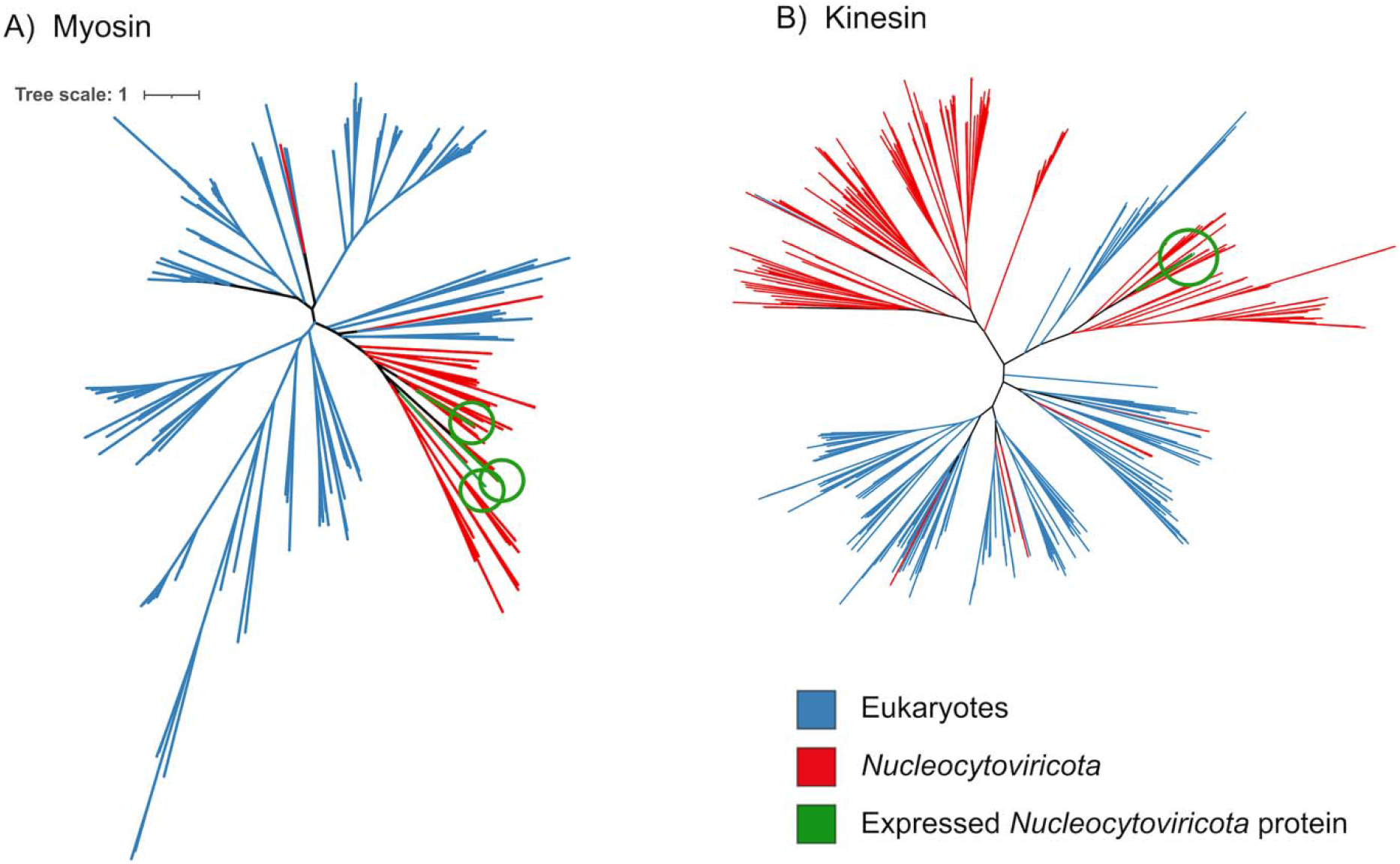
Phylogeny of *Nucleocytoviricota* myosin (left) and kinesin (right) proteins together with references available in the EggNOG database. The myosin and kinesin genes that were identified in the metatranscriptomes are colored green.

Recently it has been shown that giant viruses commonly endogenize into the genomes of their hosts, which raises the possibility that some of the transcriptional patterns we identified in this study are due to Giant Endogenous Viral Elements (GEVEs) rather than free viruses (68). Previous examination of the transcriptional landscape of GEVEs indicated that the hallmark *Nucleocytoviricota* genes involved in information processing, such as major capsid protein and DNA polymerase are typically not expressed. While numerous genes of GEVEs show varying levels of expressions, others lack detectable expression under normal growth conditions (68), contrary to patterns observed during the active viral infection (50, 52, 55). The patterns we report here for the metatranscriptomes are therefore more consistent with the gene expression patterns of viruses undergoing active viral replication rather than the transcriptional activity of GEVEs. Nonetheless, the presence of GEVEs in many algal genomes raises the possibility that not all transcription of viral genes in a microbial community can be directly linked to viral propagation.

To potentially link viruses to their hosts, we performed a network-based analysis based on correlations of whole-genome transcription of both viruses and eukaryotic plankton (Fig 6). This was motivated by several studies that have noted that the transcriptional activity of many viruses is often correlated to that of their host *in situ* (39, 49, 72). The resulting network links *Nucleocytoviricota* to diverse potential hosts, including Dinophyta, Ciliophora, Stramenopiles, and Pelagophyta. Many members of the *Prasinoviridae* clustered in the same part of the network as *Ostreococcus*, consistent with the prediction that most members of this family are Prasinoviruses. Several viruses of the family *Mesomimiviridae* were correlated with members of the diatom genera *Leptocylindrus, Chaeotoceros*, and *Eucampia*, consistent with a recent co-occurrence analysis from Tara Oceans data which suggested that some *Nucleocytoviricota* are potentially associated with diatoms (73); this association remains speculative pending more detailed analysis, however. In general, these results support the view that *Nucleocytoviricota* likely infect diverse hosts in the environment, but specific host predictions should be treated with caution pending validation with experimental approaches.

**Figure 6.**
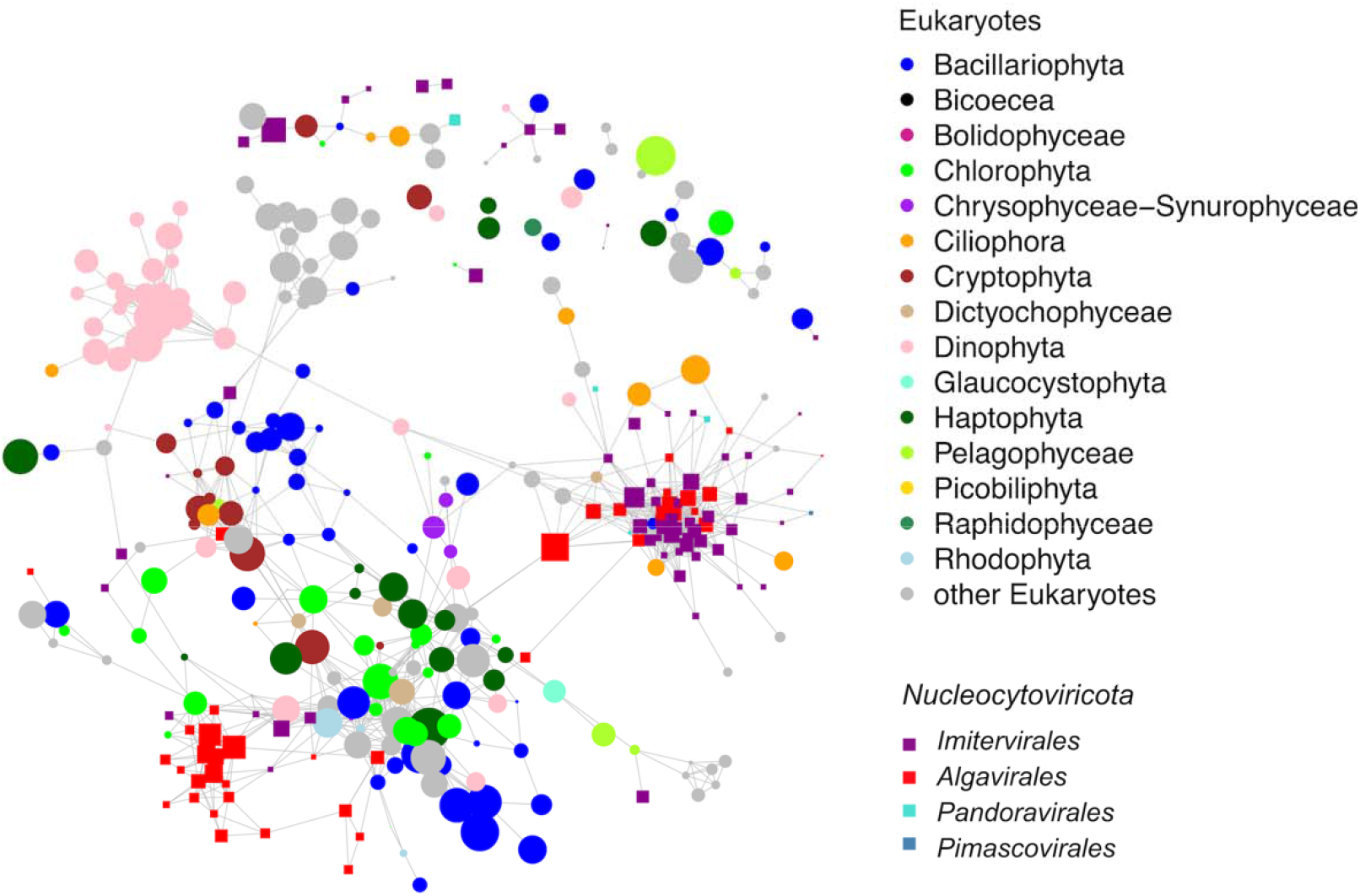
Host-virus expression correlation network. Each node represents a eukaryotic genome (circle) or viral genome (square) The line between two genomes denotes a correlation coefficient larger than 0.8. The sizes of the nodes represent the total abundance in TPM (log-transformed) of the genome and the colors show the genome’s taxonomy.

## Conclusion

In this study, we identified 145 *Nucleocytoviricota* with high transcriptional activity over a 60 hour period in surface waters of a coastal marine system. We analyzed only genomes for which > 10% of the predicted genes could be recovered at some point during the time-series; this provides high confidence that the detected viruses were indeed present, but it also implies that the actual number of active viruses is far higher than 145 because many viral transcripts remain below the level of detection. This therefore further highlights the remarkable diversity of active *Nucleocytoviricota* in this coastal system and implies more generally that these viruses are an important component of global marine environments. In addition, we found expression of diverse viral-encoded functional genes from central metabolic pathways as well as genes with similarity to eukaryotic cytoskeletal components, emphasizing the complex mechanisms employed by giant viruses to manipulate their hosts in the environment. One implication of these findings is that these viruses also have a potentially important role in shaping marine food webs; the relative impact of top-down vs bottom up controls in marine systems has long been debated (74), and the ability of *Nucleocytoviricota* to infect both heterotrophic protists (i.e., grazers) and phototrophic plankton (i.e. primary producers) suggests they can influence the relative impact of these forces. Future work identifying the hosts of these viruses and their relative contribution to eukaryotic plankton mortality rates will therefore be important avenues of future work. Overall, our findings suggest that *Nucleocytoviricota* are an underappreciated component of marine systems that exert an important influence on community dynamics and biogeochemical cycling.

## Methods

### Compilation of a *Nucleocytoviricota* Genome Database for Read Mapping

We compiled a set of *Nucleocytoviricota* genomes for transcript mapping that included metagenome-assembled genomes (MAGs) as well as genomes of cultured isolates. For this we first downloaded 2,074 genomes from Schulz et al. (70) and 501 genomes from Moniruzzaman et al. (12), two recent studies which generated numerous *Nucleocytoviricota* MAGs. We also included all *Nucleocytoviricota* genomes available in NCBI RefSeq as of June 1st, 2020. Lastly, we also included several *Nucleocytoviricota* genomes from select publications that were not yet available in NCBI, such as the cPacV, ChoanoV, and AbALV viruses that have recently been described (43, 50, 75).

After compiling this set, we dereplicated the genomes, since the presence of highly similar or identical genomes can obfuscate the results of read mapping. For dereplication we compared all *Nucleocytoviricota* genomes against each other using MASH v. 2.0 (76) (“mash dist” parameters -k 16 and -s 300), and clustered genomes together using a single-linkage clustering, with all genomes with a MASH distance of <= 0.05 linked together. The MASH distance of 0.05 was chosen since it has been found that it corresponds to an average nucleotide identity (ANI) of 95% (76), which has furthermore been suggested to be a useful threshold for distinguishing viral species-like clusters (77). From each cluster, we chose the genome with the highest N50 contig length as the representative. Prior to read mapping we also decontaminated the genomes through analysis with ViralRecall v.2.o (78) (-c parameter), with all contigs with negative scores removed. The final database used for read mapping contained 2,431 viral genomes.

Our read mapping strategy employed a translated homology search (DNA searched against protein), and so we predicted proteins from all genomes using Prodigal 2.6.3 (79) (run independently on each genome using default parameters), and then masked the protein database using tantan v. 22 (80) (default parameters) to prevent spurious read mapping to low complexity sequences. We then formatted the database for read mapping using the lastdb utility in LAST v. 1060 (81).

### Metatranscriptome Dataset

We examined the metatranscriptomic dataset of 16 timepoints obtained from >5 μm size fraction microbial communities previously reported for the California current system (39). In this study community RNA samples were collected every four hours over a 60-hour time frame, starting 6 AM September 16 to 6 PM September 18. Details regarding sample processing have been previously described (39, 82). We downloaded and trimmed reads from each of the 16 metatranscriptome libraries with Trim Galore v. 0.6.4 with parameters “--length 50 -q 5 – stringency 1”, and after removing singleton reads we then mapped paired reads onto the *Nucleocytoviricota* database using LASTAL v. 1060 (81) (parameters -m 10 -u 2 -Q 1 -F 15). We only considered hits with percent identity > 95% and bit score > 50. Transcript mapping workflows similar to this have been successfully employed in other studies (44, 46). For relative abundance comparisons read mapping counts were converted to Transcripts per Kilobase per Million (TPM) (83). To avoid the spurious detection of viral genomes in the metatranscriptomes we only considered genomes for which ≥ 10% of the proteins in that genome had hits; this cutoff was guided by recent work which has suggested that reads mapping to > 10% of a genome are unlikely to be due to spurious mappings, and therefore are indicative of the presence of that virus in a given sample (84). After performing this filtering, we arrived at a set of 145 *Nucleocytoviricota* reference genomes that were used in subsequent analysis.

### Protein Annotations

We annotated proteins in our *Nucleocytoviricota* database by comparing them to the EggNOG 4.5 (85) and Pfam v. 31 (86) Hidden Markov Models (HMMs) using HMMER3 3.3 (parameter “-E 1e-3” for the NOG search and “--cut_nc” for the Pfam search). In both cases only best hits were retained. These annotations are available in Supplemental Data 2. Annotations were manually reviewed to examine functions of interest, including energy metabolism, carbohydrate metabolism, cytoskeleton, membrane transport, light harvesting, and oxidative stress regulation. Hallmark *Nucleocytoviricota* proteins were identified using HMMs previously described (12).

### *Nucleocytoviricota* Phylogeny and Clade Delineation

To provide phylogenetic context for the 145 genomes we identified in the metatranscriptomes, and to delineate broader clades, we constructed a large mutli-locus phylogenetic tree of *Nucleocytoviricota* using the marker genes DNA polymerase B (PolB), A32 packaging enzyme (A32), Superfamily II helicase (SFII), VLTF3 transcription factor (VLTF3), RNA Polymerase large subunit (RNAPL), Topoisomerase family II (TopoII), and transcription factor IIB (TFIIB). We previously benchmarked this marker set and found that it is suitable for making phylogenetic trees across all 6 *Nucleocytoviricota* orders (5). For this tree we used all 145 genomes present in the metatranscriptomes as well as select reference *Nucleocytoviricota* sampled from across all of the main orders (*Asfuvirales, Chitovirales, Pimascovirales, Algavirales, Pandoravirales*, and *Imitervirales*). The concatenated phylogeny was generated with the ncldv_markersearch.py script, with default parameters (github.com/faylward/ncldv_markersearch), the alignment was trimmed with TrimAl v. 1.4.rev22 (87) (parameter -gt 0.1), and the tree was inferred using IQ-TREE (88) with the LG+F+I+G4 model. Support values were inferred using 1000 ultrafast bootstraps (89). We previously delineated family-level and subfamily-level clades for the *Nucleocytoviricota* (12), and we used this nomenclature to do the same for this tree.

### Virus-Host Transcriptional Network

To identify potential marine host-virus relationships, we computed a Pearson correlation matrix between the total abundance of all genomes with gene expression across the 16 timepoints (function cor() in R). The total abundance of genomes was calculated as the summed abundance of all genes of the genome expressed. The abundance of individual genes was calculated by normalizing the number of raw reads by the total number of raw reads per million in each sample. Only correlations coefficients between genome pairs exceeding 0.8 were reported.

### Tests for Diel Cycling and Day/Night Overexpression

To explore the transcriptional activities and possible diel expression pattern of the *Nucleocytoviricota* community, we examined the TPM-normalized transcriptional profiles of individual *Nucleocytoviricota* genes using RAIN (90). We did the same for each of the 145 *Nucleocytoviricota* genomes present in the transcriptomes by summing TPM expression values across each genome. P-values for these tests were corrected using the Benjamini and Hochberg method (91) as implemented by the p.adjust command in R.

In addition to strict tests for diel periodicity, we also sought to examine potential overexpression of *Nucleocytoviricota* in daytime vs nighttime periods. For this we performed a Mann-Whitney U-test on the summed gene expression of each virus, using timepoints of 0600, 1000, and 1400 as daytime and timepoints of 1800, 2200, and 0200 as nighttime (wilcox.test in R). For these tests, we also corrected the p-values using the same method as the RAIN tests.

To find clusters (modules) of genomes with highly correlated temporal expression, we performed the Weighted Gene Co-expression Network Analysis method (92) on the TPM-normalized transcriptional profiles of the *Nucleocytoviricota* genome-wide transcriptional profiles. We used a soft thresholding power of 14, parameters TOMType =“signed”, reassignThreshold = 0, minModuleSize=10 and mergeCutHeight = 0.25. We then used the moduleEigengenes utility in the WGCNA package to summarize the genome module’s average expression profile (module eigengene).

### Phylogeny of Myosin and Kinesin Proteins

To examine the evolutionary history of the *Nucleocytoviricota* proteins with myosin and kinesin domains we identified all occurrences of these proteins in our viral genome database, including those proteins that were not expressed in the metatranscriptomes. For this we only considered proteins with matches to the PF00063 (myosin) and PF00225 (kinesin) (only scores greater than the noise cutoff considered). For references we downloaded all the kinesin (COG5059) and myosin (COG5022) homologs from the EggNOG database v4.5. Many of the species harbor multiple of these proteins, and for the purpose of broad phylogenetic analysis, we randomly selected one copy of these genes per species. Diagnostic phylogenetic trees were constructed using FastTree v. 2 (93) implemented in ete3 toolkit (94) for the purpose of removing long branches and redundant or small sequences (< 300 amino acids long). To construct the final trees, we used ClustalOmega to align the sequences and trimAl (parameter -gt 0.1) to trim the alignments. We used IQ-TREE to build maximum likelihood phylogenies with the model LG+I+G4 and assessed the node support with 1000 ultrafast bootstrap replicates.

## Data Availability

The datasets analyzed in this study are already publicly available and were accessed as described in the Methods section.

## Supplemental Documents

**Supplemental Figure S1.** Histogram of genome size distribution of the 145 *Nucleocytoviricota* viruses present in the metatranscriptomes.

**Supplemental Figure S2.** *Nucleocytoviricota* metabolic gene expression in each of the 16 timepoints. The y-axis shows the functional annotation; the x-axis denotes the timepoints with the numbers after the underscores indicating the hours. The sizes of the bubbles represent the total abundance in TPM (log-transformed) of the gene across all 16 timepoints and the colors show the functional categories. Abbreviations: PFK: Phosphofructokinase, PGM: Phosphoglycerate mutase, GAPDH: Glyceraldehyde 3-P dehydrogenase, PEPCK: Phosphoenolpyruvate carboxykinase, FBP1: Fructose-1-6-bisphosphatase, SDH: Succinate dehydrogenase, 6PGD: 6-phosphogluconate dehydrogenase, LHCB: Chlorophyll a/b binding protein, ACAD: Acyl-CoA dehydrogenase, HAT: Histone acetyltransferase, GST: Glutathione S-transferase.

**Supplemental Data S1**. Summary of expression level of the 145 viruses with mapping in the metatranscriptomes. Column name abbreviations: *num_genes_expressed:* the number of genes in the genome that were expressed in at least one transcriptome;*sum_TPM, mean_TPM, median TPM:* the total, average and median abundances (in TPM) of genomes across 16 timepoints, respectively; *num_timepoints_expressed:* number of timepoints at which the gene was expressed; *proteins*: the total number of protein encoded in the viral genome; *percent_proteins*: the percentage of proteins in the genome that had hits in at least one metatranscriptome; *NCLDV_markers:* the presence or absence of the four marker genes A32, major capsid protein (mcp), SFII, and PolB in the genome; *mwt_day_night_pval:* p-values of Mann-Whitney U test for significantly different expression during daytime versus nighttime periods; *mwt_diff:* the difference between the total expression during daytime versus nighttime periods; *overexpressed*: the time period when the genome is significantly higher expressed according to Mann-Whitney U test; *Cor_Euk:* names of eukaryotic species that have significant correlation in expression (i.e. correlations coefficients exceeding 0.8) with the virus; *Correlation_strength:*correlations coefficient values corresponding to the Eukaryotes listed in the *Cor_Euk* column.

**Supplemental Data S2**. Information on *Nucleocytoviricota* genes expressed in the transcriptomes: Total expression across all 16 timepoints (*Sum_TPM*), number of timepoints at which the gene was expressed (*num_timepoints_expressed*), gene annotations and similarity bit score with reference to the databases EggNOG 4.5 and Pfam v. 31, and marker gene annotations (*NCLDV_marker*).

## Acknowledgments

We acknowledge the use of the Virginia Tech Advanced Research Computing Center for bioinformatic analyses performed in this study. We are thankful to the members of Aylward Lab and Andrew Allen for their help with a previous version of this manuscript.

## Notes

### Competing Interest Statement

The authors have declared no competing interest.

### Summary of Updates

Some references have been fixed, and some additional text has been added to the Discussion. The nomenclature used for the taxonomy has also been changed.

## References

1. Fischer MG. 2016. Giant viruses come of age. Curr Opin Microbiol 31:50–57.

2. Koonin EV. 2005. Virology: Gulliver among the Lilliputians. Current Biology 15:R167–R169.

3. Koonin EV, Yutin N. 2010. Origin and evolution of eukaryotic large nucleo-cytoplasmic DNA viruses. Intervirology 53:284–292.

4. Koonin EV, Dolja VV, Krupovic M, Varsani A, Wolf YI, Yutin N, Zerbini FM, Kuhn JH. 2020. Global organization and proposed megataxonomy of the virus world. Microbiol Mol Biol Rev 84:e00061–19.

5. Aylward FO, Moniruzzaman M, Ha AD, Koonin EV. 2021. A phylogenomic framework for charting the diversity and evolution of giant viruses. bioRxiv 20210505442809.

6. Koonin EV, Yutin N. 2019. Chapter Five - Evolution of the large nucleocytoplasmic DNA viruses of Eukaryotes and convergent origins of viral gigantism, p. 167–202. In Advances in Virus Research. Academic Press.

7. Karki S, Moniruzzaman M, Aylward FO. 2021. Comparative genomics and environmental distribution of large dsDNA viruses in the family Asfarviridae. Front Microbiol 12:657471.

8. Weynberg KD, Allen MJ, Wilson WH. 2017. Marine Prasinoviruses and their tiny plankton hosts: A review. Viruses 9:43.

9. Abergel C, Legendre M, Claverie J-M. 2015. The rapidly expanding universe of giant viruses: Mimivirus, Pandoravirus, Pithovirus and Mollivirus. FEMS Microbiol Rev 39:779–796.

10. Claverie J-M, Abergel C. 2018. Mimiviridae: An expanding family of highly diverse large dsDNA viruses infecting a wide phylogenetic range of aquatic eukaryotes. Viruses 10:506.

11. Blanc-Mathieu R, Dahle H, Hofgaard A, Brandt D, Ban H, Kalinowski J, Ogata H, Sandaa R-A. 2021. A persistent giant algal virus, with a unique morphology, encodes an unprecedented number of genes involved in energy metabolism. J Virol 95:e02446–20.

12. Moniruzzaman M, Martinez-Gutierrez CA, Weinheimer AR, Aylward FO. 2020. Dynamic genome evolution and complex virocell metabolism of globally-distributed giant viruses. Nat Commun 11:1710.

13. Rodrigues RAL, Arantes TS, Oliveira GP, Silva LKS, Abrahão JS. 2019. The complex nature of Tupanviruses, p. 135–166. In Advances in Virus Research. Academic Press.

14. Moreau H, Piganeau G, Desdevises Y, Cooke R, Derelle E, Grimsley N. 2010. Marine prasinovirus genomes show low evolutionary divergence and acquisition of protein metabolism genes by horizontal gene transfer. J Virol 84:12555–12563.

15. Yutin N, Koonin EV. 2012. Proteorhodopsin genes in giant viruses. Biol Direct 7:34.

16. Rozenberg A, Oppermann J, Wietek J, Fernandez Lahore RG, Sandaa R-A, Bratbak G, Hegemann P, Béjà O. 2020. Lateral gene transfer of anion-conducting channelrhodopsins between green algae and giant viruses. Curr Biol 30:4910–4920.e5.

17. Wilson WH, Schroeder DC, Allen MJ, Holden MTG, Parkhill J, Barrell BG, Churcher C, Hamlin N, Mungall K, Norbertczak H, Quail MA, Price C, Rabbinowitsch E, Walker D, Craigon M, Roy D, Ghazal P. 2005. Complete genome sequence and lytic phase transcription profile of a Coccolithovirus. Science 309:1090–1092.

18. Monier A, Pagarete A, de Vargas C, Allen MJ, Read B, Claverie J-M, Ogata H. 2009. Horizontal gene transfer of an entire metabolic pathway between a eukaryotic alga and its DNA virus. Genome Res 19:1441–1449.

19. Schvarcz CR, Steward GF. 2018. A giant virus infecting green algae encodes key fermentation genes. Virology 518:423–433.

20. Forterre P. 2013. The virocell concept and environmental microbiology. ISME J 7:233–236.

21. Forterre P. 2011. Manipulation of cellular syntheses and the nature of viruses: The virocell concept. C R Chim 14:392–399.

22. Rosenwasser S, Ziv C, van Creveld SG, Vardi A. 2016. Virocell metabolism: Metabolic innovations during host-virus interactions in the ocean. Trends Microbiol 24:821–832.

23. Ogata H, Toyoda K, Tomaru Y, Nakayama N, Shirai Y, Claverie J-M, Nagasaki K. 2009. Remarkable sequence similarity between the dinoflagellate-infecting marine girus and the terrestrial pathogen African swine fever virus. Virol J 6:178.

24. Sandaa RA, Heldal M, Castberg T, Thyrhaug R, Bratbak G. 2001. Isolation and characterization of two viruses with large genome size infecting Chrysochromulina ericina (Prymnesiophyceae) and Pyramimonas orientalis (Prasinophyceae). Virology 290:272–280.

25. Brussaard CPD, Short SM, Frederickson CM, Suttle CA. 2004. Isolation and phylogenetic analysis of novel viruses infecting the phytoplankton Phaeocystis globosa (Prymnesiophyceae). Appl Environ Microbiol 70:3700–3705.

26. Moniruzzaman M, LeCleir GR, Brown CM, Gobler CJ, Bidle KD, Wilson WH, Wilhelm SW. 2014. Genome of brown tide virus (AaV), the little giant of the Megaviridae, elucidates NCLDV genome expansion and host-virus coevolution. Virology 466-467:60–70.

27. Ghedin E, Claverie J-M. 2005. Mimivirus relatives in the Sargasso sea. Virol J 2:62.

28. Monier A, Larsen JB, Sandaa R-A, Bratbak G, Claverie J-M, Ogata H. 2008. Marine mimivirus relatives are probably large algal viruses. Virol J 5:12.

29. Williamson SJ, Rusch DB, Yooseph S, Halpern AL, Heidelberg KB, Glass JI, Andrews-Pfannkoch C, Fadrosh D, Miller CS, Sutton G, Frazier M, Venter JC. 2008. The Sorcerer II Global Ocean Sampling expedition: Metagenomic characterization of viruses within aquatic microbial samples. PLoS One 3:e1456.

30. Chen F, Suttle CA, Short SM. 1996. Genetic diversity in marine algal virus communities as revealed by sequence analysis of DNA polymerase genes. Appl Environ Microbiol 62:2869–2874.

31. Endo H, Blanc-Mathieu R, Li Y, Salazar G, Henry N, Labadie K, de Vargas C, Sullivan MB, Bowler C, Wincker P, Karp-Boss L, Sunagawa S, Ogata H. 2020. Biogeography of marine giant viruses reveals their interplay with eukaryotes and ecological functions. Nat Ecol Evol 4:1639–1649.

32. Kaneko H, Blanc-Mathieu R, Endo H, Chaffron S, Delmont TO, Gaia M, Henry N, Hernández-Velázquez R, Nguyen CH, Mamitsuka H, Forterre P, Jaillon O, de Vargas C, Sullivan MB, Suttle CA, Guidi L, Ogata H. 2021. Eukaryotic virus composition can predict the efficiency of carbon export in the global ocean. iScience 24:102002.

33. Mihara T, Koyano H, Hingamp P, Grimsley N, Goto S, Ogata H. 2018. Taxon richness of “Megaviridae” exceeds those of bacteria and archaea in the ocean. Microbes and Environments 33:162–171.

34. Li Y, Hingamp P, Watai H, Endo H, Yoshida T, Ogata H. 2018. Degenerate PCR primers to reveal the diversity of giant viruses in coastal waters. Viruses 10:496.

35. Laber CP, Hunter JE, Carvalho F, Collins JR, Hunter EJ, Schieler BM, Boss E, More K, Frada M, Thamatrakoln K, Brown CM, Haramaty L, Ossolinski J, Fredricks H, Nissimov JI, Vandzura R, Sheyn U, Lehahn Y, Chant RJ, Martins AM, Coolen MJL, Vardi A, DiTullio GR, Van Mooy BAS, Bidle KD. 2018. Coccolithovirus facilitation of carbon export in the North Atlantic. Nat Microbiol 3:537–547.

36. Sheyn U, Rosenwasser S, Lehahn Y, Barak-Gavish N, Rotkopf R, Bidle KD, Koren I, Schatz D, Vardi A. 2018. Expression profiling of host and virus during a coccolithophore bloom provides insights into the role of viral infection in promoting carbon export. ISME J 12:704–713.

37. Zeigler Allen L, McCrow JP, Ininbergs K, Dupont CL, Badger JH, Hoffman JM, Ekman M, Allen AE, Bergman B, Venter JC. 2017. The Baltic sea virome: Diversity and transcriptional activity of DNA and RNA viruses. mSystems 2:e00125–16.

38. Carradec Q, Pelletier E, Da Silva C, Alberti A, Seeleuthner Y, Blanc-Mathieu R, Lima-Mendez G, Rocha F, Tirichine L, Labadie K, Kirilovsky A, Bertrand A, Engelen S, Madoui M-A, Méheust R, Poulain J, Romac S, Richter DJ, Yoshikawa G, Dimier C, Kandels-Lewis S, Picheral M, Searson S, Tara Oceans Coordinators, Jaillon O, Aury J-M, Karsenti E, Sullivan MB, Sunagawa S, Bork P, Not F, Hingamp P, Raes J, Guidi L, Ogata H, de Vargas C, Iudicone D, Bowler C, Wincker P. 2018. A global ocean atlas of eukaryotic genes. Nat Commun 9:373.

39. Kolody BC, McCrow JP, Allen LZ, Aylward FO, Fontanez KM, Moustafa A, Moniruzzaman M, Chavez FP, Scholin CA, Allen EE, Worden AZ, Delong EF, Allen AE. 2019. Diel transcriptional response of a California Current plankton microbiome to light, low iron, and enduring viral infection. ISME J 13:2817–2833.

40. Santini S, Jeudy S, Bartoli J, Poirot O, Lescot M, Abergel C, Barbe V, Wommack KE, Noordeloos AAM, Brussaard CPD, Claverie J-M. 2013. Genome of Phaeocystis globosa virus PgV-16T highlights the common ancestry of the largest known DNA viruses infecting eukaryotes. Proc Natl Acad Sci U S A 110:10800–10805.

41. Stough JMA, Yutin N, Chaban YV, Moniruzzaman M, Gann ER, Pound HL, Steffen MM, Black JN, Koonin EV, Wilhelm SW, Short SM. 2019. Genome and environmental activity of a Chrysochromulina parva virus and its virophages. Front Microbiol 10:703.

42. Gallot-Lavallée L, Blanc G, Claverie J-M. 2017. Comparative genomics of Chrysochromulina ericina virus and other microalga-infecting large DNA viruses highlights their intricate evolutionary relationship with the established Mimiviridae family. J Virol 91:e00230–17.

43. Needham DM, Yoshizawa S, Hosaka T, Poirier C, Choi CJ, Hehenberger E, Irwin NAT, Wilken S, Yung C-M, Bachy C, Kurihara R, Nakajima Y, Kojima K, Kimura-Someya T, Leonard G, Malmstrom RR, Mende DR, Olson DK, Sudo Y, Sudek S, Richards TA, DeLong EF, Keeling PJ, Santoro AE, Shirouzu M, Iwasaki W, Worden AZ. 2019. A distinct lineage of giant viruses brings a rhodopsin photosystem to unicellular marine predators. Proceedings of the National Academy of Sciences 116:20574–20583.

44. Aylward FO, Eppley JM, Smith JM, Chavez FP, Scholin CA, DeLong EF. 2015. Microbial community transcriptional networks are conserved in three domains at ocean basin scales. Proc Natl Acad Sci U S A 112:5443–5448.

45. Derelle E, Yau S, Moreau H, Grimsley NH. 2018. Prasinovirus attack of Ostreococcus is furtive by day but savage by night. J Virol 92:e01703–17.

46. Aylward FO, Boeuf D, Mende DR, Wood-Charlson EM, Vislova A, Eppley JM, Romano AE, DeLong EF. 2017. Diel cycling and long-term persistence of viruses in the ocean’s euphotic zone. Proc Natl Acad Sci U S A 114:11446–11451.

47. Hevroni G, Flores-Uribe J, Béjà O, Philosof A. 2020. Seasonal and diel patterns of abundance and activity of viruses in the Red Sea. Proc Natl Acad Sci U S A 117:29738–29747.

48. Yoshida T, Nishimura Y, Watai H, Haruki N, Morimoto D, Kaneko H, Honda T, Yamamoto K, Hingamp P, Sako Y, Goto S, Ogata H. 2018. Locality and diel cycling of viral production revealed by a 24 h time course cross-omics analysis in a coastal region of Japan. The ISME Journal 12:1287–1295.

49. Moniruzzaman M, Wurch LL, Alexander H, Dyhrman ST, Gobler CJ, Wilhelm SW. 2017. Virus-host relationships of marine single-celled eukaryotes resolved from metatranscriptomics. Nat Commun 8:16054.

50. Needham DM, Poirier C, Hehenberger E, Jiménez V, Swalwell JE, Santoro AE, Worden AZ. 2019. Targeted metagenomic recovery of four divergent viruses reveals shared and distinctive characteristics of giant viruses of marine eukaryotes. Philos Trans R Soc Lond B Biol Sci 374:20190086.

51. Rodrigues RAL, Louazani AC, Picorelli A, Oliveira GP, Lobo FP, Colson P, La Scola B, Abrahão JS. 2020. Analysis of a Marseillevirus transcriptome reveals temporal gene expression profile and host transcriptional shift. Front Microbiol 11:651.

52. Legendre M, Audic S, Poirot O, Hingamp P, Seltzer V, Byrne D, Lartigue A, Lescot M, Bernadac A, Poulain J, Abergel C, Claverie J-M. 2010. mRNA deep sequencing reveals 75 new genes and a complex transcriptional landscape in Mimivirus. Genome Res 20:664–674.

53. Lartigue A, Burlat B, Coutard B, Chaspoul F, Claverie J-M, Abergel C. 2015. The megavirus chilensis Cu,Zn-superoxide dismutase: the first viral structure of a typical cellular copper chaperone-independent hyperstable dimeric enzyme. J Virol 89:824–832.

54. Moniruzzaman M, Gann ER, Wilhelm SW. 2018. Infection by a giant virus (AaV) induces widespread physiological reprogramming in Aureococcus anophagefferens CCMP1984 - A harmful bloom algae. Front Microbiol 9:752.

55. Monier A, Chambouvet A, Milner DS, Attah V, Terrado R, Lovejoy C, Moreau H, Santoro AE, Derelle É, Richards TA. 2017. Host-derived viral transporter protein for nitrogen uptake in infected marine phytoplankton. Proc Natl Acad Sci U S A 114:E7489–E7498.

56. Monier A, Welsh RM, Gentemann C, Weinstock G, Sodergren E, Armbrust EV, Eisen JA, Worden AZ. 2012. Phosphate transporters in marine phytoplankton and their viruses: cross-domain commonalities in viral-host gene exchanges. Environ Microbiol 14:162–176.

57. Aherfi S, Belhaouari DB, Pinault L, Baudoin J-P, Decloquement P, Abrahao J, Colson P, Levasseur A, Lamb DC, Chabriere E, Raoult D, La Scola B. 2020. Tricarboxylic acid cycle and proton gradient in Pandoravirus massiliensis: Is it still a virus? bioRxiv 20200921306415.

58. Vincent F, Sheyn U, Porat Z, Vardi A. 2021. Visualizing active viral infection reveals diverse cell fates in synchronized algal bloom demise. Proceedings of the National Academy of Sciences 118:e2021586118.

59. Gastrich MD, Leigh-Bell JA, Gobler, CH, Anderson, OR, Wilhelm, SW, Bryan M. 2004. Viruses as potential regulators of regional brown tide blooms caused by the alga, Aureococcus anophagefferens 27:112–119.

60. Raoult D, Audic S, Robert C, Abergel C, Renesto P, Ogata H, La Scola B, Suzan M, Claverie J-M. 2004. The 1.2-megabase genome sequence of Mimivirus. Science 306:1344–1350.

61. Yoosuf N, Yutin N, Colson P, Shabalina SA, Pagnier I, Robert C, Azza S, Klose T, Wong J, Rossmann MG, La Scola B, Raoult D, Koonin EV. 2012. Related giant viruses in distant locations and different habitats: Acanthamoeba polyphaga moumouvirus represents a third lineage of the Mimiviridae that is close to the Megavirus lineage. Genome Biol Evol 4:1324–1330.

62. Silva LCF, Almeida GMF, Assis FL, Albarnaz JD, Boratto PVM, Dornas FP, Andrade KR, La Scola B, Kroon EG, da Fonseca FG, Abrahão JS. 2015. Modulation of the expression of mimivirus-encoded translation-related genes in response to nutrient availability during Acanthamoeba castellanii infection. Front Microbiol 6:539.

63. Saini HK, Fischer D. 2007. Structural and functional insights into Mimivirus ORFans. BMC Genomics 8:1–12.

64. Abrahão JS, Araújo R, Colson P, La Scola B. 2017. The analysis of translation-related gene set boosts debates around origin and evolution of mimiviruses. PLoS Genet 13:e1006532.

65. Da Cunha V, Gaia M, Ogata H, Jaillon O, Delmont TO, Forterre P. 2020. Giant viruses encode novel types of actins possibly related to the origin of eukaryotic actin: the viractins. bioRxiv 20200616150565.

66. Subramaniam K, Behringer DC, Bojko J, Yutin N, Clark AS, Bateman KS, van Aerle R, Bass D, Kerr RC, Koonin EV, Stentiford GD, Waltzek TB. 2020. A new family of DNA viruses causing disease in crustaceans from diverse aquatic biomes. MBio 11: e02938–19.

67. Kijima S, Delmont TO, Miyazaki U, Gaia M, Endo H, Ogata H. Discovery of viral myosin genes with complex evolutionary history within plankton. bioRxiv 2021.03.14.435220.

68. Moniruzzaman M, Weinheimer AR, Martinez-Gutierrez CA, Aylward FO. 2020. Widespread endogenization of giant viruses shapes genomes of green algae. Nature 588:141–145.

69. Guglielmini J, Woo AC, Krupovic M, Forterre P, Gaia M. 2019. Diversification of giant and large eukaryotic dsDNA viruses predated the origin of modern eukaryotes. Proc Natl Acad Sci U S A 116:19585–19592.

70. Schulz F, Roux S, Paez-Espino D, Jungbluth S, Walsh DA, Denef VJ, McMahon KD, Konstantinidis KT, Eloe-Fadrosh EA, Kyrpides NC, Woyke T. 2020. Giant virus diversity and host interactions through global metagenomics. Nature 578:432–436.

71. Nelson DR, Hazzouri KM, Lauersen KJ, Jaiswal A, Chaiboonchoe A, Mystikou A, Fu W, Daakour S, Dohai B, Alzahmi A, Nobles D, Hurd M, Sexton J, Preston MJ, Blanchette J, Lomas MW, Amiri KMA, Salehi-Ashtiani K. 2021. Large-scale genome sequencing reveals the driving forces of viruses in microalgal evolution. Cell Host Microbe 29:250–266.e8.

72. Stough JMA, Kolton M, Kostka JE, Weston DJ, Pelletier DA, Wilhelm SW. 2018. Diversity of active viral infections within the Sphagnum microbiome. Appl Environ Microbiol 84:e01124–18.

73. Meng L, Endo H, Blanc-Mathieu R, Chaffron S, Hernández-Velázquez R, Kaneko H, Ogata H. 2020. Quantitative assessment of NCLDV–host interactions predicted by co-occurrence analyses. bioRxiv 20201016342030.

74. Lynam CP, Llope M, Möllmann C, Helaouët P, Bayliss-Brown GA, Stenseth NC. 2017. Interaction between top-down and bottom-up control in marine food webs. Proc Natl Acad Sci U S A 114:1952–1957.

75. Matsuyama T, Takano T, Nishiki I, Fujiwara A, Kiryu I, Inada M, Sakai T, Terashima S, Matsuura Y, Isowa K, Nakayasu C. 2020. A novel Asfarvirus-like virus identified as a potential cause of mass mortality of abalone. Sci Rep 10:4620.

76. Ondov BD, Treangen TJ, Melsted P, Mallonee AB, Bergman NH, Koren S, Phillippy AM. 2016. Mash: Fast genome and metagenome distance estimation using MinHash. Genome Biol 17:132.

77. Bobay L-M, Ochman H. 2018. Biological species in the viral world. Proceedings of the National Academy of Sciences 115:6040–6045.

78. Aylward FO, Moniruzzaman M. 2021. ViralRecall-A flexible command-line tool for the detection of giant virus signatures in ‘omic data. Viruses 13:150.

79. Hyatt D, Chen G-L, Locascio PF, Land ML, Larimer FW, Hauser LJ. 2010. Prodigal: Prokaryotic gene recognition and translation initiation site identification. BMC Bioinformatics 11:119.

80. Frith MC. 2011. A new repeat-masking method enables specific detection of homologous sequences. Nucleic Acids Res 39:e23.

81. Kiełbasa SM, Wan R, Sato K, Horton P, Frith MC. 2011. Adaptive seeds tame genomic sequence comparison. Genome Res 21:487–493.

82. Shi Y, Tyson GW, DeLong EF. 2009. Metatranscriptomics reveals unique microbial small RNAs in the ocean’s water column. Nature 459:266–269.

83. Mortazavi A, Williams BA, McCue K, Schaeffer L, Wold B. 2008. Mapping and quantifying mammalian transcriptomes by RNA-Seq. Nat Methods 5:621–628.

84. Tithi SS, Aylward FO, Jensen RV, Zhang L. 2018. FastViromeExplorer: A pipeline for virus and phage identification and abundance profiling in metagenomics data. PeerJ 6:e4227.

85. Huerta-Cepas J, Szklarczyk D, Forslund K, Cook H, Heller D, Walter MC, Rattei T, Mende DR, Sunagawa S, Kuhn M, Jensen LJ, von Mering C, Bork P. 2016. eggNOG 4.5: A hierarchical orthology framework with improved functional annotations for eukaryotic, prokaryotic and viral sequences. Nucleic Acids Res 44:D286–93.

86. El-Gebali S, Mistry J, Bateman A, Eddy SR, Luciani A, Potter SC, Qureshi M, Richardson LJ, Salazar GA, Smart A, Sonnhammer ELL, Hirsh L, Paladin L, Piovesan D, Tosatto SCE, Finn RD. 2019. The Pfam protein families database in 2019. Nucleic Acids Res 47:D427–D432.

87. Capella-Gutiérrez S, Silla-Martínez JM, Gabaldón T. 2009. trimAl: A tool for automated alignment trimming in large-scale phylogenetic analyses. Bioinformatics 25:1972–1973.

88. Nguyen L-T, Schmidt HA, von Haeseler A, Minh BQ. 2015. IQ-TREE: A fast and effective stochastic algorithm for estimating maximum-likelihood phylogenies. Mol Biol Evol 32:268–274.

89. Hoang DT, Chernomor O, von Haeseler A, Minh BQ, Vinh LS. 2018. UFBoot2: Improving the ultrafast bootstrap approximation. Mol Biol Evol 35:518–522.

90. Thaben PF, Westermark PO. 2014. Detecting rhythms in time series with RAIN. J Biol Rhythms 29:391–400.

91. Benjamini Y, Hochberg Y. 1995. Controlling the false discovery rate: A practical and powerful approach to multiple testing. Journal of the Royal Statistical Society: Series B (Methodological) 57:289–300.

92. Langfelder P, Horvath S. 2008. WGCNA: an R package for weighted correlation network analysis. BMC Bioinformatics 9:1–13.

93. Price MN, Dehal PS, Arkin AP. 2010. FastTree 2--approximately maximum-likelihood trees for large alignments. PLoS One 5:e9490.

94. Huerta-Cepas J, Dopazo J, Gabaldón T. 2010. ETE: a Python environment for tree exploration. BMC Bioinformatics 11:24.

